# Metabolic dysfunction promoted by mitochondrial DNA mutation burden drives retinal degeneration

**DOI:** 10.1101/2025.07.30.667736

**Authors:** Johnathon Sturgis, Ke Jiang, Stephanie A. Hagstrom, Maya Moorthy, Vera L. Bonilha

## Abstract

Retinal degenerative diseases, such as age-related macular degeneration (AMD), retinitis pigmentosa, and glaucoma, have been linked to mitochondrial dysfunction. However, the impact of mitochondrial DNA (mtDNA) mutation accumulation in the context of these retinopathies has yet to be thoroughly explored. Our previous studies focused on the retinal phenotype observed in the PolgD257A mutator mice (D257A), revealing the effects of aging and mtDNA mutation accumulation in the retina. We have reported that this model exhibited significant morphological and functional deficits in the retina by 6 months of age, with notable alterations in the retinal pigment epithelium (RPE) occurring as early as 3 months, including changes in the cristae density and reduction in length of mitochondria.

This study investigated how mtDNA mutations affect the metabolic interaction between the retina and RPE in young (3 months) and old (12 months) wild-type (WT) and D257A mice. We assessed cellular energy production using freshly dissected retina samples from both groups through Seahorse analysis, immunofluorescence, and Western blot experiments.

The analysis of aged D257A retina punches revealed significantly reduced basal and maximal mitochondrial respiration, along with increased mitochondrial reserve capacity compared to WT. However, glycolytic flux, measured as a function of extracellular acidification rate (ECAR), did not differ between WT and D257A mice. Both D257A retina and RPE exhibited decreased expression of essential electron transport proteins involved in oxidative phosphorylation. Additionally, we observed a reduction in the expression of glucose transporter 1 (GLUT-1) and lactate transporter (MCT1) at the apical surface of the RPE. Enzymes associated with glycolysis, including hexokinase II and lactate dehydrogenase A, were significantly lower in the aged D257A retina, while hexokinase I and pyruvate kinase 2 were upregulated in the RPE.

These findings indicate that the accumulation of mtDNA mutations leads to impaired metabolism in both the retina and RPE. Furthermore, it suggests that glucose from the choroidal blood supply is being utilized by the RPE rather than being transported to the neural retina. Mitochondrial dysfunction in RPE promotes a glycolytic state in these cells, leading to reduced availability of metabolites and, consequently, diminished overall retinal function. These results are essential for advancing our understanding of the mechanisms underlying retinal degeneration and provide a new perspective on the role of mtDNA mutations in these diseases.

## Introduction

Retinal degeneration is a clinically relevant biological phenomenon associated with numerous age-related eye disorders, including diabetic retinopathy, glaucoma, retinitis pigmentosa, and age-related macular degeneration (AMD) ^[1–3]^ It is widely believed that retinal degeneration originates from an energetic imbalance between the photoreceptors (PRs) and the retinal pigment epithelium (RPE), resulting in photoreceptor cell death, RPE atrophy, inflammation, and ultimately, loss of vision ^[4]^. The challenge in investigating degenerative retinal diseases lies in understanding the molecular mechanisms of early disease stages and their progression to vision loss. Currently, it is believed that multiple factors contribute to disease etiology, including genetic and environmental risks, inflammation, reactive oxygen species (ROS), and, more recently, mitochondrial dysfunction ^[5]^. More specifically, mitochondrial DNA (mtDNA) damage has been implicated in the development of several ocular pathologies including AMD, retinitis pigmentosa, optic atrophy, and Leber’s hereditary optic neuropathy ^[6, 7]^. Furthermore, studies analyzing human donor tissue detected significant mtDNA damage in the RPE of donors affected by late stages of AMD ^[8]^.

An additional factor to consider in understanding the role of mtDNA mutation accumulation in these age-related ocular diseases is the contribution of aging. As we all age, we accumulate mutations in our mitochondrial genome, which primarily impact post-mitotic tissues such as the retina that do not have a source of cell renewal. The accumulation of these mutations is hypothesized to be accelerated in disease conditions ^[9]^. The prevailing notion for decades has been that mtDNA mutations arise from a “vicious cycle theory”, in which mtDNA mutations from ROS-mediated damage lead to increased ROS production and further mtDNA damage or mutation accumulation ^[10]^. However, recent advances in sequencing technology have suggested that this phenomenon may not be the primary cause of aging and age-associated diseases. A more recently proposed hypothesis of aging suggests that mtDNA mutations arise from errors in mtDNA replication and repair mechanisms, and these mutations clonally expand as we age ^[11]^. Once these mutations reach a certain threshold, overall mitochondrial function in a cell is compromised. Over time, most mitochondria in the tissue become dysfunctional, leading to progressive degeneration ^[12]^.

Here, we investigated the consequences of mtDNA mutation accumulation on retinal metabolism in the PolgD257A (D257A) mouse. This mouse model has an aspartic acid to alanine substitution at the 257th amino acid residue of the exonuclease domain of the Polymerase gamma (PolG) gene; the protein responsible for mtDNA replication, proofreading, and repair ^[13]^. This mutation renders PolG incapable of mtDNA repair, resulting in an mtDNA mutation accumulation rate 2,500 times higher than that of the wild-type mice. Our initial characterization of this mouse model revealed key insights into when mtDNA mutation accumulation starts to impact the function and morphology of the retina. Importantly, this study did not reveal increases in oxidative stress associated with mtDNA mutation burden in the retina or RPE. Instead, our study identified the downregulation of essential electron transport chain proteins at early time points in this model, which may promote an energy deficit in cells crucial for retinal function. Finally, changes in mitochondrial-specific proteins in the retina, as well as alterations in RPE mitochondrial morphology, were observed ^[14]^.

Studies from several labs have demonstrated that the retina and RPE work together as a metabolic ecosystem ^[15]^. The RPE relies on reductive carboxylation and oxidative phosphorylation for energy production, whereas the photoreceptor cells in the outer retina rely on glycolysis ^[16, 17]^. Thus, dysfunctional RPE mitochondria upset the established balance between RPE and outer retinal metabolism. Therefore, understanding the energetic balance of the retina is crucial for understanding the pathology and treatment of retinal diseases ^[18]^. We believe that mitochondrial dysfunction, driven by the accumulation of mtDNA mutations, alters the metabolic activity of retinal cells, leading to degeneration over time. Our study adopts a novel approach to this issue, focusing on the role of mtDNA mutations in retinal degeneration. Our research lays the groundwork for further exploration of these hypotheses and their potential implications for understanding and treating retinal degeneration. As such, the current study aims to elucidate the impact of the mtDNA mutation burden on retina and RPE metabolism during aging. We use diverse and corroborative methodologies to understand how mitochondrial and non-mitochondrial-associated metabolic pathways are affected by increased mtDNA mutational burden.

## Results

### Decreased oxygen consumption in D257A mice retina

Our previous retinal characterization of the D257A mouse model detected electron transport chain (ETC) protein deficiencies as early as 3 and 6 months of age ^[14]^. Interestingly, the proteins found to be significantly depleted in this study were complex I (CI) and complex IV (CIV), both of which contain multiple subunits encoded by mtDNA. These findings were the first indication of potential respiratory chain dysfunction in the D257A retina and RPE. To determine if this dysfunction altered mitochondrial activity, we measured the oxygen consumption rate (OCR) of D257A mice’ ex-vivo retinal punches using a previously established protocol ^[19]^. Using the well-established Seahorse Bioanalyzer, we performed a mitochondria stress test and measured both basal and maximal OCR in 3- and 12-month WT and D257A mice retinal punches (Fig. 1A). Basal respiration in 12-month D257A mice retina was significantly decreased compared to 12- month WT mice (Fig. 1B). After injection of Bam15, a reagent that uncouples mitochondrial respiration from the proton gradient, maximal respiration was determined. Maximal respiration was significantly decreased in 3-month D257A retina compared to 3-month WT and was more significantly decreased in 12-month D257A retina compared to 12-month WT (Fig. 1C). Next, mitochondrial reserve capacity (MRC), which represents the difference between the cell’s normal energy production and its maximum energy production from oxygen consumption, was determined by subtracting the calculated basal respiration rate from the maximal respiration rate and then dividing by the basal respiration rate, expressed as a percentage. MRC was significantly increased in 12-month D257A retina compared to 12-month WT (Fig. 1D). To determine if these results were due to the increased retinal degeneration previously observed in the aging D257A mouse retina and therefore indicative of less respiration due to the loss of retinal cells rather than metabolic perturbation, protein concentration was determined in 3- and 12-month WT and D257A retinal punches. Protein concentration was not significantly decreased in 3- or 12-month D257A retinas compared to WT (Fig. S1A). After the injection of rotenone and antimycin A, reagents that block electron transfer from complex I and III, respectively, and shut down oxidative phosphorylation-coupled oxygen consumption, non-mitochondrial oxygen consumption can be determined. By determining non-mitochondrial oxygen consumption, we can infer that the differences observed were specifically due to mitochondria-associated oxygen consumption. Both basal and maximal mitochondrial-associated OCR were significantly reduced in 12- month D257A retinas compared to WT (Fig. S1B, C). Together, these data confirm that the previously observed depletion of ETC proteins results in a decreased OCR in the D257A retina. Interestingly, depletion of ETC proteins precedes the loss of OCR, given that the most significant alterations to OCR were found in the 12-month D257A retina. However, maximal respiration is already altered in the 3-month D257A retina, indicating dysfunctional oxygen consumption even when this process is uncoupled from the proton gradient. Importantly, these findings are not a result of less cellular content but rather a consequence of decreased cellular activity.

**FIGURE 1:**
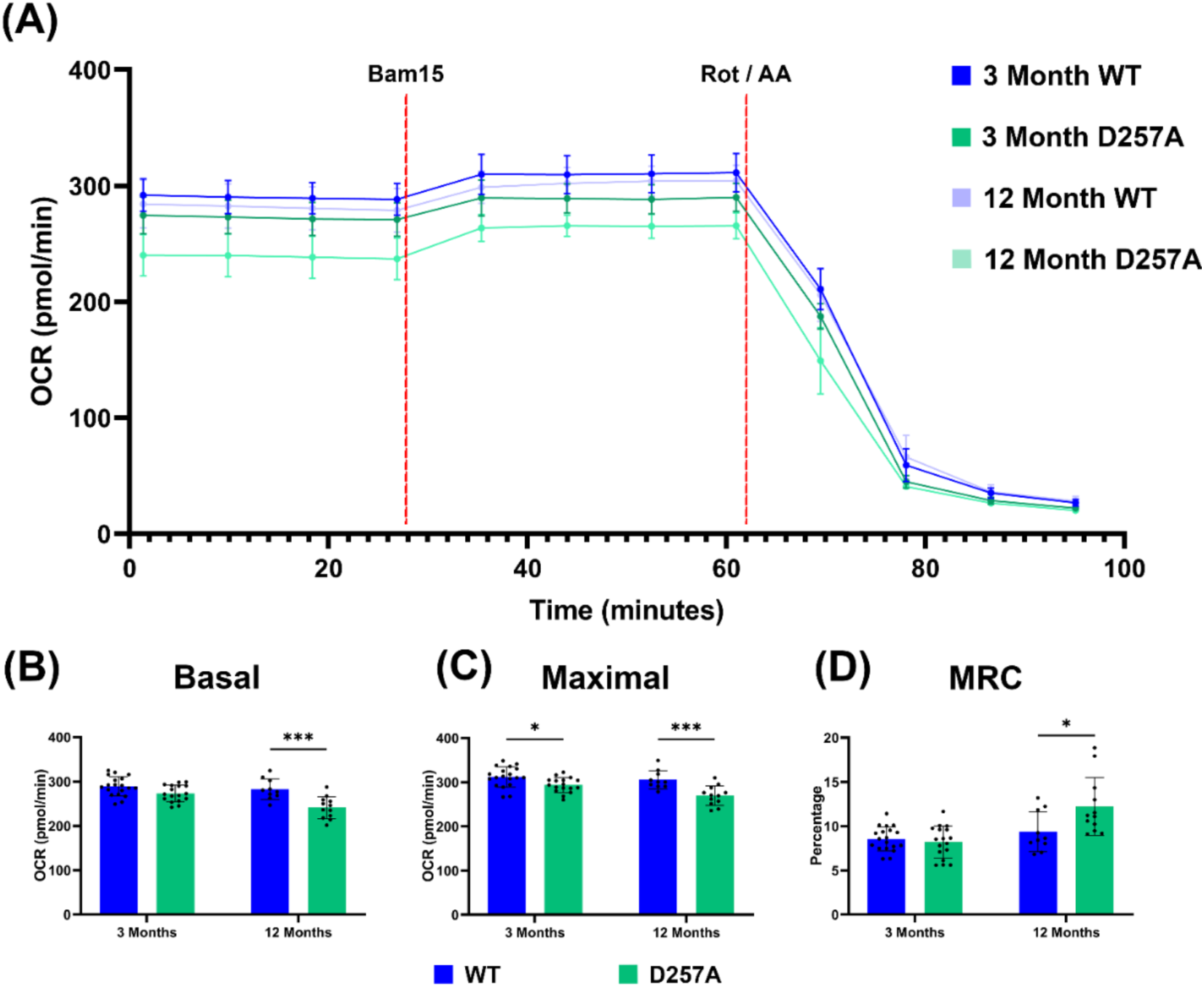
Oxygen consumption rate of neural retina punches measured by seahorse analysis. **A)** Graphical representation of seahorse program protocol. Graphical representation of **B)** Basal respiration, **C)** Maximal respiration, and **D)** Mitochondrial reserve capacity (MRC). Data points = technical replicates / individual retina punches. Statistics: two-way ANOVA, p-value < 0.05.

### Decreased glycolysis in aged mice retina

Glycolysis is a metabolic pathway that converts glucose to pyruvate and subsequently pyruvate to lactate. When pyruvate is converted to lactate, this process releases protons that acidify the surrounding environment. For this reason, measuring extracellular acidification rate (ECAR) can give a readout for glycolysis occurring in each tissue or cell sample. We also analyzed the same retinal punches from the mice described above to measure ECAR using the Seahorse Bioanalyzer. By employing an assay strategy similar to the one used to measure OCR, we were able to determine both basal and maximal ECAR (Fig. 2A). Basal ECAR was not significantly reduced when comparing 3- or 12-month D257A retina to WT. However, basal ECAR was significantly reduced with aging in both the WT and D257A retina (Fig. 2B). After the injection of rotenone and antimycin A, the retinal cells are driven into a glycolytic state for energy production, as they can no longer receive energy from ETC-associated oxygen consumption. This treatment pushes the tissue towards a glycolytic state and allows us to determine maximal ECAR. Like basal ECAR levels, maximal ECAR was not significantly altered when comparing 3- or 12-month D257A retina to WT. However, maximal ECAR was significantly reduced with aging in both the WT and D257A retina (Fig. 2C). Next, glycolytic reserve capacity (GRC), a representation of the difference between the cell’s normal energy production and its maximum energy production from glycolysis, was determined by subtracting the calculated basal ECAR from the maximal ECAR and dividing by basal ECAR to be represented as a percentage. GRC was not significantly altered when comparing 3- or 12-month D257A retina to WT. However, GRC was significantly increased with aging in both the WT and D257A retina (Fig. 2D). After injection of 2-Deoxy-D-glucose (2-DG), a glucose analog that inhibits glycolysis, non-glycolytic ECAR can be determined. By determining non-glycolytic ECAR, we can infer that the differences observed were specifically due to glycolysis associated with ECAR. Both basal and maximal glycolytic-associated ECAR were significantly decreased with age in 12-month WT and D257A retina (Fig. 2B, C). Together, these results indicate that mtDNA mutation accumulation does not affect glycolytic function or capacity in the D257A mouse retina. However, between 3 and 12 months of age, the mouse retina decreases its glycolysis based on ECAR measurements. Our data suggest that age-related decreases in glycolysis could be indicative of glucose-deficiency or decreased glucose related metabolism in the neural retina. Additionally, the decrease in glycolysis could be due to a decrease in the number of rod and cone photoreceptors with aging. This phenomenon could be also attributed to the RPE cells using the glucose as an energy source instead.

**FIGURE 2:**
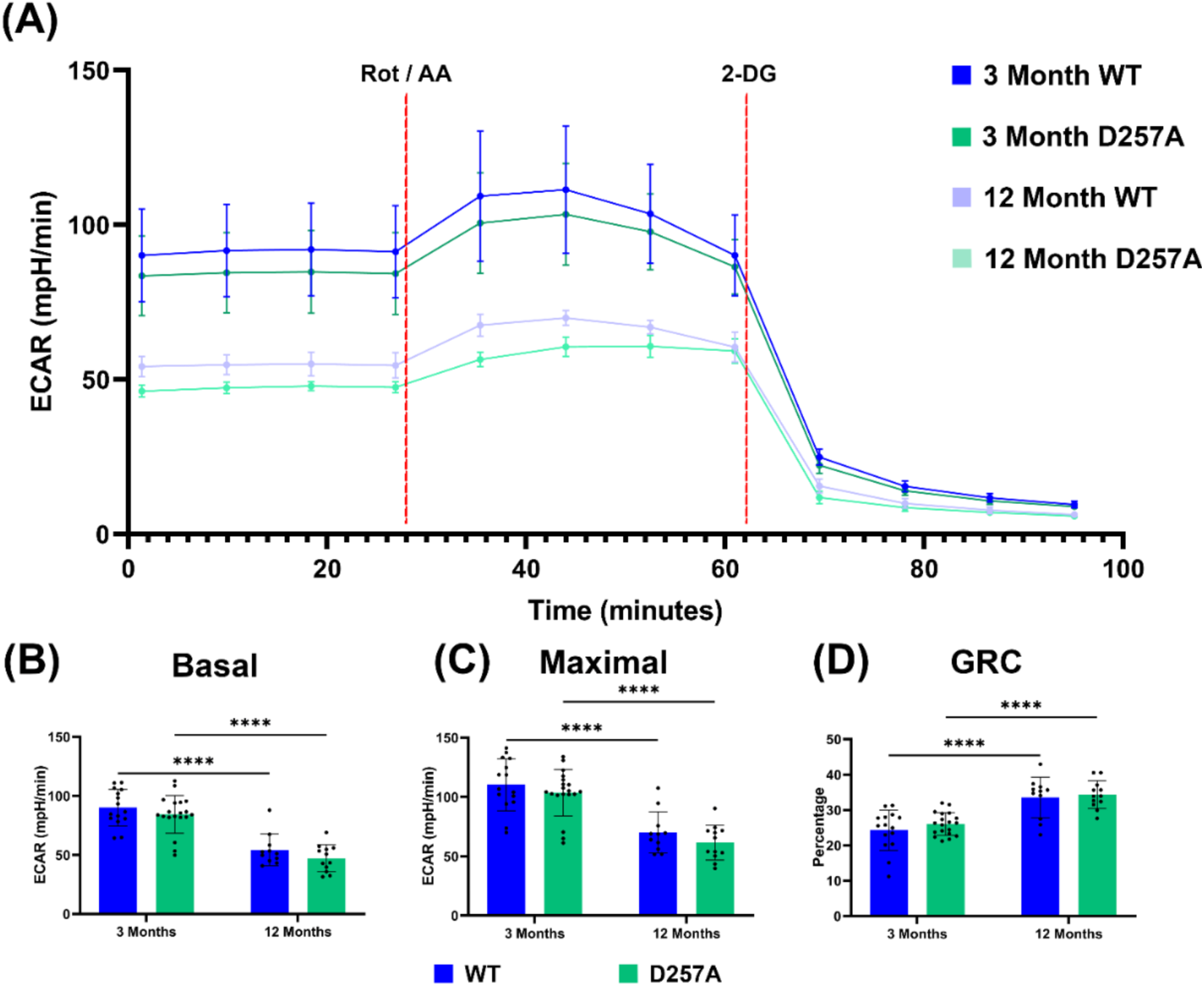
Extracellular acidification rate of neural retina punches measured by seahorse analysis. **A)** Graphical representation of seahorse program protocol. Graphical representation of **B)** Basal glycolysis, **C)** Maximal glycolysis, and **D)** Glycolytic reserve capacity (GRC). Data points = technical replicates / individual retina punches. Statistics: two-way ANOVA, p-value < 0.05.

### Electron transport chain alteration in D257A retina and RPE

Increased mtDNA mutation accumulation with age predominantly affects mtDNA-encoded proteins. These include essential subunits of the ETC required for proper assembly of the complex protein involved in oxidative phosphorylation. Previous studies on mtDNA in the mutator (D257A) mice have shown that these complexes are severely downregulated due to the mtDNA mutation burden ^[14, 20, 21]^. To determine the impact of aging and mtDNA mutation accumulation in the mitochondrial metabolism of the retina and RPE, we performed western blot analysis using an antibody cocktail probing for markers of each complex of the ETC (Fig. 3A). First, using whole retina protein lysate we determined that the levels of complex V (CV) were upregulated in the 12-month D257A compared to 12-month WT (Fig. 3B), levels of complex III (CIII) were not significantly changed at either age between WT or D257A (Fig. 3C), levels of complex IV (CIV) were significantly decreased at 3 months of age between WT and D257A but did not remain significantly decreased at 12-months of age (Fig. 3D), levels of complex II (CII) were upregulated in 12-month D257A compared to WT (Fig. 3E), and levels of complex I (CI) were significantly decreased at both 3- and 12-months of age in D257A retina (Fig. 3F). Next, we probed the same ETC components described above in RPE lysates (Fig. 4A). Quantification of RPE protein lysate signals determined that CV, CIII, and CII protein levels were not significantly altered in 3- or 12-month D257A compared to WT (Fig. 4 A, B, C, E). However, the levels of CIV were significantly decreased in both the 3- and 12-month D257A groups compared to the WT (Fig. 4D). Finally, the levels of CI were significantly decreased in 3-month D257A compared to WT (Fig. 4F). Our previous characterization of the same ETC subunits in 3- and 6-month D257A retina and RPE lysates showed similar decreases observed in this study ^[14]^. CI and CIV levels are significantly depleted in 3-month retina and RPE lysates in both studies. However, the results presented here provide more evidence on the impact of aging by probing 12-month WT and D257A retina and RPE. These complexes also appear to be the most impacted by the normal aging process. This data validates our in vivo results, again indicating that mtDNA mutation accumulation decreases the levels of proteins involved in mitochondrial-associated metabolism in both the retina and RPE of the D257A mouse.

**FIGURE 3:**
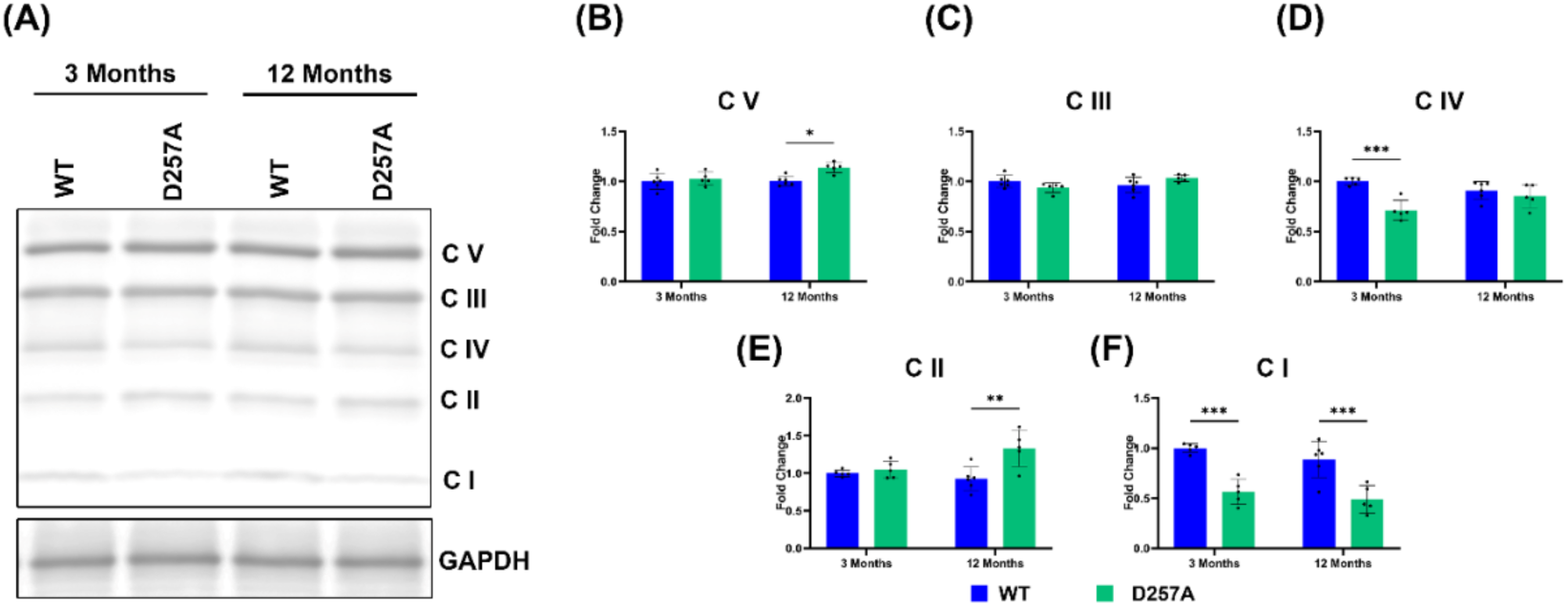
Alterations in ETC complex proteins in D257A retina. **A)** Representatives immunoblot of OXPHOS protein content in the retina. Graphical representation of **B)** CV, **C)** CIII, **D)** CIV, **E)** CII, **F)** CI Statistics: Data points = biological replicates; two-way ANOVA; p < 0.05.

**FIGURE 4:**
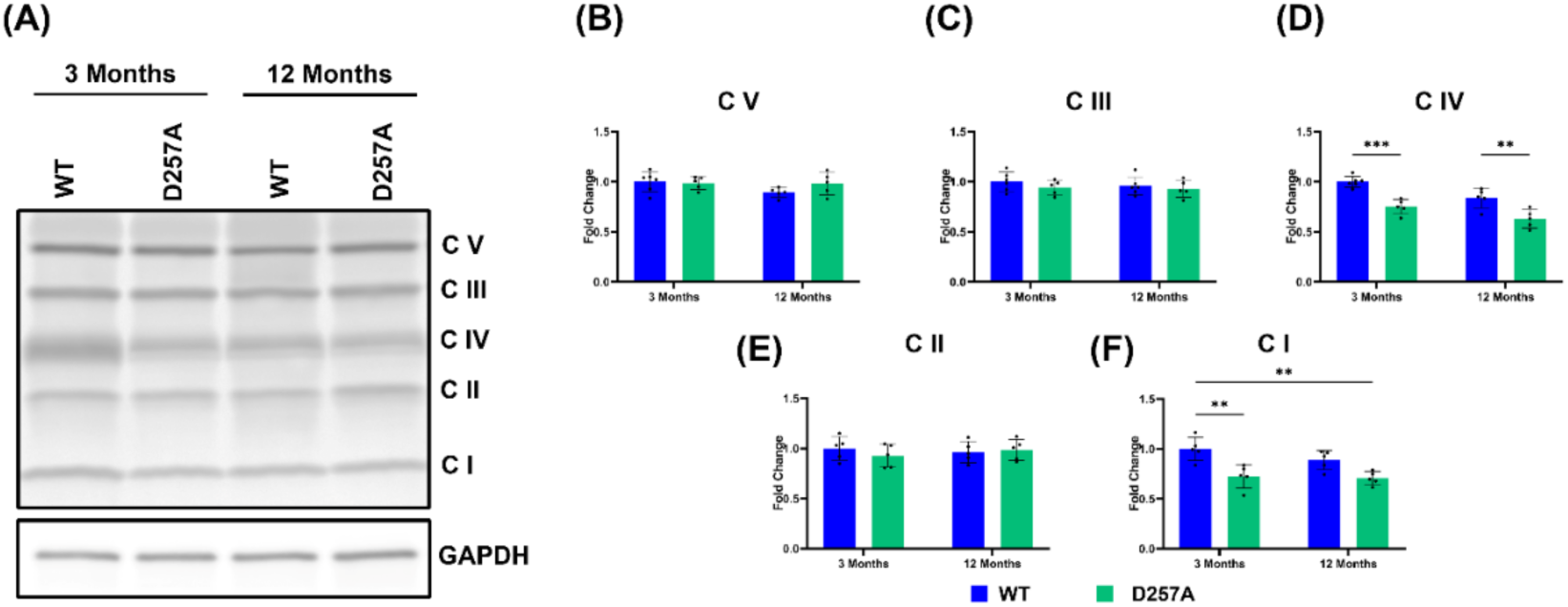
Alterations in ETC complex proteins in D257A RPE. **A)** Representatives immunoblot of OXPHOS protein content in the retina. Graphical representation of **B)** CV, **C)** CIII, **D)** CIV, **E)** CII, **F)** CI Statistics: Data points = biological replicates; two-way ANOVA; p < 0.05.

### Glycolytic enzyme protein levels in D257A retina and RPE

Given the previous evidence that mtDNA mutation accumulation results in decreased oxygen consumption, likely promoted by complex protein deficiency, we next investigated glycolysis in more detail. Our ECAR data indicated a reduction in aging-associated glycolysis but showed no evidence of a difference between the WT and D257A mice’ retinas. To further determine if glycolysis is impacted in the D257A model, we next probed retina and RPE protein lysates for various enzymes involved in the glycolytic pathway. Hexokinase (HK), the enzyme that irreversibly phosphorylates glucose in the first step of glycolysis, was investigated first. HK has two main isoforms found in the retina: HK I, which is expressed in all retinal layers, including the RPE, and HK II, which is primarily expressed in the photoreceptor cells ^[22]^. Retina lysates probed for HK II showed a significant decrease in expression of this isoform at 3 and 12 months of age in D257A mice (Fig. 5A). Retina lysates probed for HK I show no significant changes in expression of this isoform at either age in D257A mice (Fig. S2). In contrast, RPE lysates probed for HK I expression show a significant increase in this enzyme at 12 months of age in D257A mice (Fig. 5D). Next, we investigated pyruvate kinase (PKM2) expression in the retina and RPE. This enzyme catalyzes the final step in the energy-generating process of the glycolytic pathway, converting phosphoenolpyruvate (PEP) to pyruvate. Retina lysates showed no significant difference in PKM2 levels in 3- or 12-month-old D257A mice (Fig. 5B). In contrast, RPE lysates probed for this protein showed a significant increase in expression in 12-month-old D257A mice (Fig. 5E).

**FIGURE 5:**
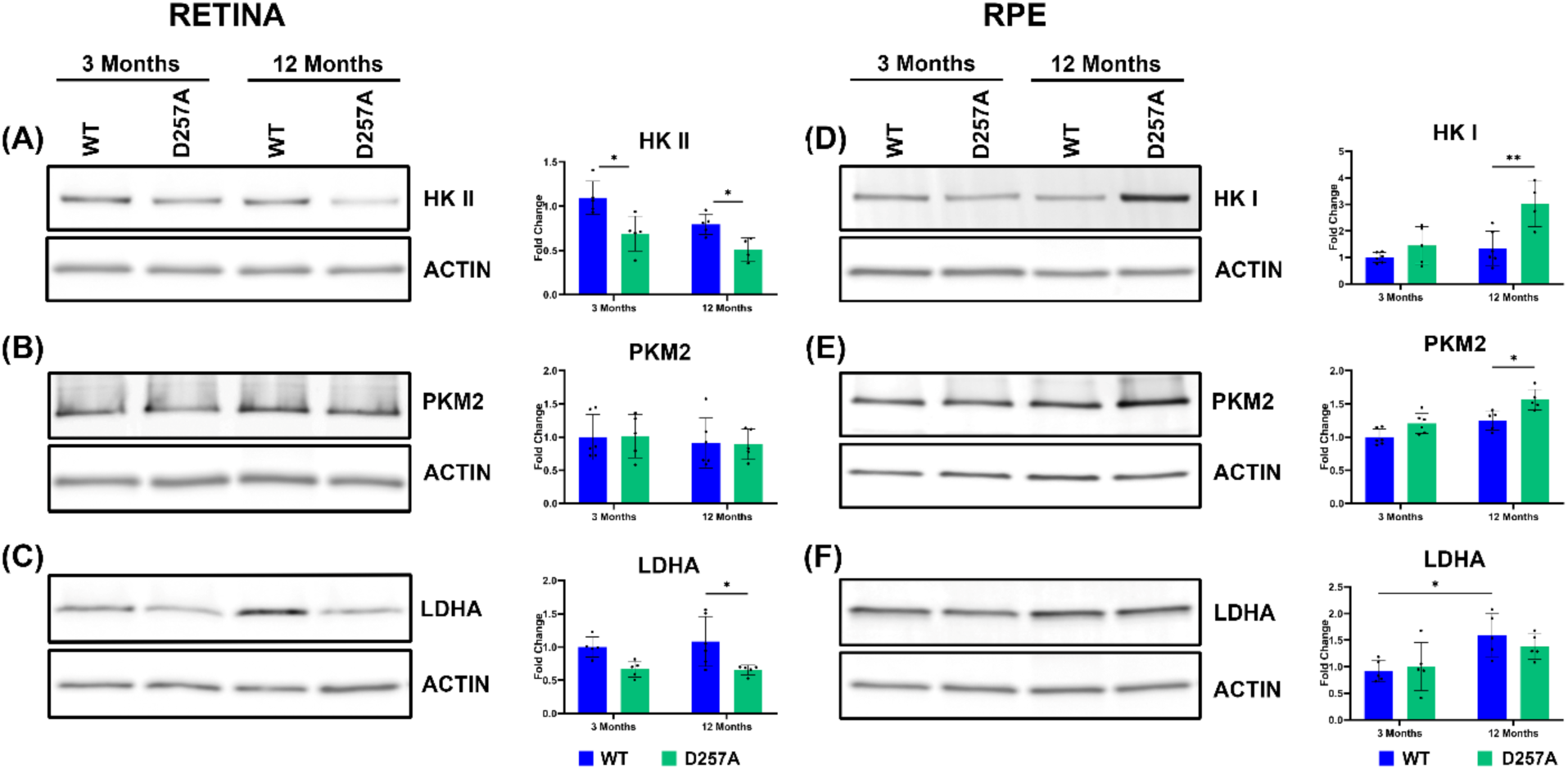
Alterations in key metabolic enzyme proteins in D257A retina and RPE. Representatives immunoblot and graphical representation of **A)** Hexokinase I (HK I), **B)** Pyruvate kinase (PKM2) **C)** Lactate dehydrogenase (LDH) in WT and D257A mouse retina. Representatives immunoblot and graphical representation of **D)** Hexokinase I (HK I),**E)** Pyruvate kinase (PKM2), **F)** Lactate dehydrogenase (LDH) in WT and D257A mouse RPE. Statistics: Data points = biological replicates (n=5); two-way ANOVA; p < 0.05.

Finally, we investigated lactate dehydrogenase (LDH) expression in the retina and RPE. LDH is expressed in two main isoforms in the retina, LDHA, which preferentially facilitates the conversion of pyruvate to lactate, and LDHB, which preferentially facilitates the conversion of lactate to pyruvate ^[23, 24]^. Retina lysates probed for the LDHA isoform showed a significant decrease in expression in 12-month D257A mice (Fig. 5C). RPE lysates probed for the same protein showed no significant difference in expression between WT and D257A mice at either time point; however, there was a significant increase in expression levels between 3-month and 12-month WT RPE (Fig. 5F). Together, these data indicate a metabolic switch occurring in the retina and RPE due to mtDNA mutation burden.

### Changes in abundance and distribution of metabolite transporter proteins in D257A RPE

Given our previous examination of altered glycolytic enzyme expression in the retina and RPE, we next aimed to conclude this study by investigating the expression levels and distribution of specific metabolite transporter protein in this model. The canonical pathway of outer retinal metabolism is that glucose is taken up from the choroidal blood supply by the basal expression of glucose transporter 1 (GLUT-1) on RPE cells. This process occurs in the basal infoldings of the RPE, which provide the necessary surface area for nutrient uptake and absorption. Glucose is then transferred to the neural retina through apical RPE expression of this same transporter. Retinal GLUT-1 then transports this glucose from the extracellular space into cells, where it is utilized for energy production. In the outer retina, the primary cells that utilize this glucose are the photoreceptors ^[17]^. The lactate produced as a byproduct of glycolysis in these cells is then transported out of the photoreceptors and taken up by the RPE by the apical lactate transporter monocarboxylate transporter 1 (MCT1) ^[25]^. Excess lactate taken up by the RPE can be returned to circulation by monocarboxylate transporter 3 (MCT3), which is expressed on the basal side, near the choroidal blood flow ^[25]^. To determine if GLUT-1 expression is altered in D257A retina, we probed retinal cryosections for this transporter protein. No significant difference was observed in retinal GLUT-1 distribution between ages or between WT and D257A mice (Fig. S3A, B). However, RPE GLUT-1 distribution was significantly decreased in 3- and 12-month-old WT mice. Interestingly, this decrease was not observed between 3- and 12-month D257A mice, which showed a significant increase in GLUT-1 distribution between 12-month WT and D257A RPE (Fig. 6A, B). Next, we probed sections for MCT1 and observed a significantly decreased RPE MCT1 distribution in WT and D257A mice between 3 and 12 months (Fig. 6A, C). Finally, we investigated MCT3 expression in RPE and found no significant difference in this transporter distribution with age or between WT and D257A mice (Fig. 6A, D).

**FIGURE 6:**
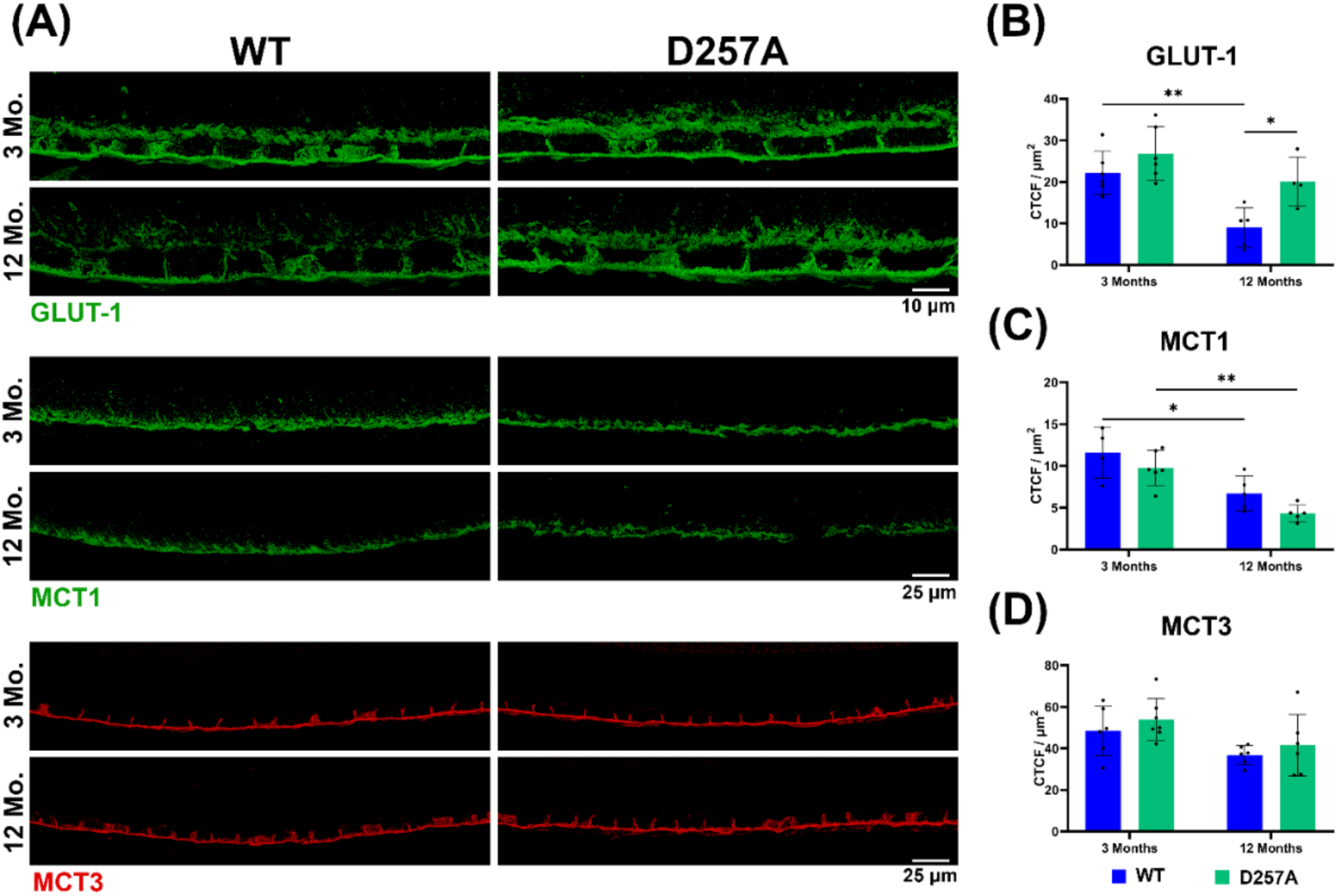
Investigation of transporters distribution and signal intensity in D257A RPE. **A)** Representative confocal images of 3-month and 12-month-old RPE metabolite transporter proteins. Graphical representation of **B)** Glucose transporter type 1 (GLUT-1), **C)** Monocarboxylate transporter 1 (MCT1), and **D)** Monocarboxylate transporter 3 (MCT3) levels in WT and D257A RPE. Statistics: Data points = biological replicates (n=5-7); two-way ANOVA; p < 0.05.

Together, these findings further suggest a glycolytic shift in RPE metabolism, corroborating the enzymatic data presented earlier. The decreases in GLUT-1 distribution observed in aged WT RPE suggest that aging alone may decrease glucose availability in the retina, possibly due to reduced glycolysis, as shown in our Seahorse ECAR data. To compensate for reduced glucose availability, the RPE may rely on other metabolites, such as glutamine. When probing for expression of the glutamate, aspartate, and cysteine excitatory amino acid transporter 3 (EAAT3) in RPE protein lysates, we found a significant increase in this protein’s expression between 3- and 12- month WT mice (Fig. S3C, D). The significant increase in GLUT-1 expression in 12- month D257A RPE compared to WT is likely due to these cells relying predominantly on glucose as their primary fuel source.

## Discussion

This study builds on the initial retinal characterization of the D257A mouse model for studying retinal-related health and degeneration to elucidate the impact of mtDNA mutation burden on retina and RPE metabolism. Total intracellular ATP is generated by two energetic pathways, glycolysis and mitochondrial OXPHOS. Alterations of mtDNA due to dysfunctional Polg induce loss of mitochondrial OXPHOS and mitochondrial ATP generation with compensatory upregulation of the cytoplasmic glycolysis process, thus increasing the contribution of glycolysis to the total cellular ATP generation ^[26]^.

Seahorse analysis of oxygen consumption rate in WT vs. D257A mouse retina shows that mutation burden results in significantly decreased basal and maximal respiration, with a significant increase in MRC. Extracellular acidification rate as a measurable function of glycolysis shows significant decreases with age in both WT and D257A retina. Thus, glycolysis in the D257A retina may be maintained by compensatory metabolism in other retinal cells that produce intermediate glycolytic byproducts. Additional experiments in the retinas of WT and D257A mice would be necessary to understand better why glycolysis is not altered in these mice.

Electron transport chain proteins show alterations in both the retina and RPE of D257A mice. Interestingly, proteins most encoded by mtDNA seem to be preferentially depleted (CIV and CI). These results confirm data presented in our initial characterization of this model and provide further evidence of the implications for age-associated mtDNA mutation burden. In addition, some complex proteins (CV and CII), encoded by nuclear DNA, appear to be upregulated in the retinas of the D257A model. The change in expression of these ETC proteins could be due to mitochondrial-nuclear crosstalk, in which cells sense dysfunctional mitochondria and attempt to compensate for this defect by translating more mitochondrial-associated proteins ^[27]^. Regardless, mtDNA mutation accumulation decreases the levels of proteins involved in mitochondrial-associated metabolism in both the retina and RPE of the D257A mouse. This accumulation of mtDNA mutations will likely have a direct impact on the transcription and translation of ETC complexes. However, another aspect of the ETC complexes is their specific activity. Our experiments did not investigate the assembly of functional ETC complexes. Future experiments on ETC complex-specific activity are needed to address this question. Previous reports in skeletal muscle ETC complex specific activity did not detect significant differences between genotypes in any of the complex activities, suggesting that either only functional complexes assemble, or dysfunctional complexes are effectively degraded ^[28]^.

The most intriguing evidence for a glycolytic shift occurring in the outer retina is that the enzymes involved in glycolysis are altered from their normal state. Photoreceptor glycolysis seems to be specifically decreased in the neural retina due to the loss of HK II expression ^[29]^. Previous studies of this protein have also demonstrated its necessity for maintaining photoreceptor survival during aging and outer retinal stress ^[29]^. The significant increase in HK I and PKM2 expression in the D257A RPE also indicates that these cells are becoming more glycolytic. HK mediates the initial enzymatic reaction in glycolysis that converts glucose to glucose-6-phosphate I in cells. Importantly, this is an irreversible reaction, essentially trapping glucose inside the cell. The increase in PKM2 provides further evidence that the glycolytic process is occurring in the RPE of the aged D257A mice. A previous study reported that Polg binds to PKM2 and affects the activation of its Tyr105-site phosphorylation, thus interfering with the glycolysis of gastric cancer cells ^[30]^. Our results agree with previous studies, which reported that increased glycolysis in the skeletal muscle of D257A mice was associated with higher levels of 6- phosphofructo-2-kinase/fructose-2,6-bisphosphatase 3 (PFKFB3) ^[31]^. Additionally, in lung cancer cells, mtDNA mutations increased glycolysis and led to a dependence on glucose, at the expense of a decreased NAD+/NADH ratio, which inhibited de novo serine synthesis ^[32]^.

Interestingly, LDHA expression was significantly increased in WT RPE during aging. This increase in expression could be due to the reduced lactate available for the RPE to consume from the aging WT retina, which also displayed decreased basal and maximal glycolysis. Under these conditions, WT aged RPE would increase LDHA levels to compensate for the reduction of lactate produced. As stated before, LDHA preferentially converts pyruvate to lactate ^[23]^. This increase in aged WT RPE suggests that pyruvate from either the choroid or other retinal cells is being converted into lactate, possibly indicating a shift in the normal metabolic process that occurs with aging alone. Given the evidence of increased glycolytic enzyme expression of HK I and PKM2 in the aged D257A RPE, LDHA expression was expected to be increased in these mice’s RPE as well; however, this was not the case. Perhaps these RPE have already shifted to using glucose from the choroidal blood flow, and therefore, are not starved of lactate production and do not need to increase the expression of this enzyme. In the aged D257A retina, LDHA expression was significantly decreased in comparison to WT retinas. The LDHA isoform is abundant in photoreceptors ^[33–35]^. Previous reports have shown that reduced expression of LDHA in rods diminishes the amount of lactate in the mouse retina and causes rods to be shorter than usual ^[35]^. Further experiments should quantify the length of rod photoreceptors in 12-month-old WT and D257A mice.

Our investigation into transporter proteins in the retina and RPE also suggested key insights into altered metabolism in the D257A model. GLUT-1 expression in the neural retina remains constant, indicating that this transporter is not affected by aging or the burden of mtDNA mutation. This could be due to the presence of this transporter being critical for the survival retinal cells ^[36]^. A recent work knocking down GLUT-1 in the RPE showed that the decreased transport of glucose into the outer retina in this model decreased retinal glucose levels, affected outer segment renewal and photoreceptor cell survival, and resulted in activation of Müller glial cells ^[37]^. This protein may also already be expressed to its maximum capability in the retina, meaning that a lack of glucose supply from the outer retina’s blood supply does not drive compensation through higher expression of this transporter. Interestingly, GLUT-1 expression is significantly decreased in 12-month WT RPE. Our Seahorse data indicate that glycolysis is significantly decreased with age in the neural retina. Perhaps this is because the aged RPE in the D257A model is more reliant on glucose itself and therefore maintains expression of this transporter to supply itself with an alternative fuel source. Alternatively, the mitochondrial dysfunction in the D257A RPE may have resulted in slow β-oxidation and caused lipid accumulation. Thus, it could force the RPE to consume more glucose as previously reported ^[38, 39]^.

The decreased MCT1 expression on the apical surface of the 12-month WT and D257A RPE is likely driven by decreased lactate availability from the neural retina. To compensate for the loss of lactate availability, the 12-month WT RPE increases expression of glutamate transporter protein EAAT3. However, expression of this protein is not significantly increased in 12-month D257A RPE, indicating that loss of lactate supply from the neural retina does not stimulate glutamate uptake in this model. The primary glutamate transporter expressed in the photoreceptors and bipolar cells is excitatory amino acid transporter 2 and 5 (EAAT2 and EAAT5) ^[40]^. We did not investigate the expression levels of these proteins were not investigated in the current study.

In summary, this study provides sufficient evidence that mtDNA mutation burden has a significant impact on outer retinal metabolism. Previous studies have identified outer retinal metabolism as crucial for survival and maintenance of photoreceptor cells and retinal health ^[17]^. The long-standing notion of RPE and photoreceptor metabolism has been that glucose from the choroidal blood flow is taken up by the RPE and passed to the photoreceptors for use in glycolysis. The lactate produced as a byproduct of glycolysis is then exported out of the photoreceptors and taken up by the RPE. The RPE then converts this lactate into pyruvate, which is used in the TCA cycle and subsequently in oxidative phosphorylation. This pathway appears to remain constant in young (3-month) WT and D257A retina and RPE. However, in the old (12-month) D257A model, this process is disrupted. Glucose from the choroidal blood flow appears to be utilized by the RPE cells themselves, and therefore not transported to the photoreceptors. This results in a significant increase in expression of GLUT-1 and a decrease in MCT1 in the aged D257A RPE. The result from this metabolic shift that occurs with aging and mtDNA mutation burden is a degenerating retina at the cost of RPE cell survival (Fig. 7). The healthy function of the RPE is critical to maintain neural retina integrity and this has been furthermore highlighted in this study. A previous study using an RPE-selective model with complete loss of mtDNA transcription and replication results in RPE dedifferentiation, hypertrophy, and activation of mTOR which consequently drives photoreceptor degeneration^[41]^. Our data indicate that aging alone also has an impact on outer retinal metabolism. However, mtDNA mutation burden has a more severe impact on overall retinal health decline. The accumulation of mtDNA mutations promotes mitochondrial dysfunction, leading to cells becoming less dependent on mitochondrial-related metabolic pathways. Future studies should examine metabolite deficiencies and additional sources of energy available to the RPE in this model.

**FIGURE 7:**
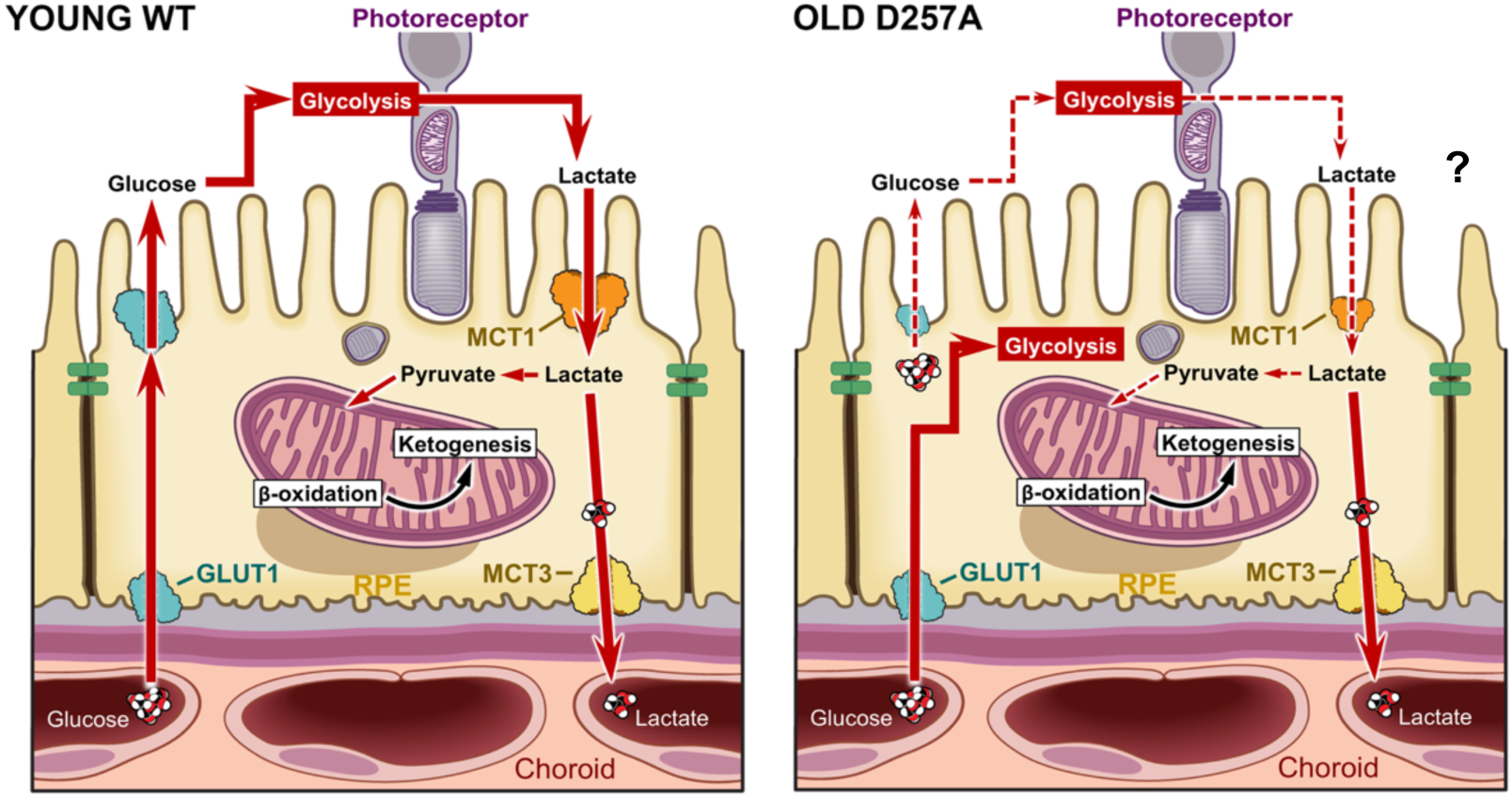
Graphical summary of key metabolic pathways found to be disrupted in D257A mice.

The findings presented here have some limitations regarding the methodology. First, the retina explants used in this study contain the entire neural retina; therefore, we cannot conclude that there are photoreceptor-specific deficits. Secondly, although the reduction in oxygen consumption is statistically significant, it still shows a relatively mild decrease. This observation may be related to the heterogeneity of the retinal tissue and the fact that one specific cell type may consume less oxygen than other cell types. Additionally, the significant reduction in glycolysis observed with aging in the retina remains to be investigated. Our previous characterization of this model indicated a significant reduction in essential rod and cone photoreceptor proteins rhodopsin and red/green opsin ^[14]^. For this reason, the reduction in glycolysis observed could be driven by loss of photoreceptor cell viability. Third, it is worth noting that inferring results from the detection of enzymatic protein expression and distribution alone is insufficient to understand the entirety of these cells’ metabolism. A more effective methodology for gaining insight into the metabolic changes occurring in this model would be to perform targeted or untargeted metabolomics, or to measure the levels of metabolite byproducts, such as lactate and glucose, in the retina and RPE separately. Finally, we did not measure the activity of the ETC complexes and transporters.

## Methods

### Mice

All procedures were approved by the Institutional Animal Care and Use Committee (ARC 2021-2191 and 003449) of the Cleveland Clinic. Twelve-week-old D257A and WT mice were purchased from the Jackson Laboratory. Experiments were conducted on male and female littermate Polg^WT/WT^ (wild-type, WT) and Polg^D257A/D257A^ (D257A). Mice were genotyped as previously described ^[42]^. To not introduce mtDNA mutation burden during birth and development, male heterozygous D257A mice were crossed with C57Bl/6J females to generate heterozygous; then male and female D257A/WT were bred to generate WT, D257A/WT and homozygous D257A/D257A experimental mice. Mice were housed in individually ventilated cages in a 14-h light/10-h dark cycle and were provided regular chow and water ad libitum. Mice retina and RPE tissue were collected at ∼ 3 hours after light input to avoid circadian variations.

### Seahorse analysis

Seahorse analysis was conducted on freshly dissected ex-vivo retinal punches to determine oxygen consumption rate (OCR) and extracellular acidification rate (ECAR) using the Seahorse XFe24 Islet Capture FluxPak (Agilent Technologies, Inc, Wilmington, DE, USA, 103518-100), as previously described ^[19]^. One day prior to the experiment, sensor cartridges were soaked in calibration solution overnight at RT. The Seahorse DMEM assay medium containing 6 mM glucose, 0.12 mM of pyruvate, and 0.5 mM of glutamine was prepared fresh the day of the experiment. Mesh inserts in the islet capture microplate were coated with the Corning Cell-Tak cell attachment medium (Corning, NY, USA, 354240) according to manufacturer’s instructions to improve tissue attachment. Retinas were dissected in assay medium, and a 1mm diameter biopsy puncher was then used to take five separate retinal punches from each retina. Punches were then placed on mesh inserts and loaded into the islet capture microplate containing 500 μL assay medium. For mitochondria stress assay, punches were incubated with 5 μM of Bam15 (TimTec LLC, Tampa, FL, USA, ST056388) and 1 μM of each rotenone (Sigma-Aldrich, St. Louis, MO, USA, R8875)/antimycin A (AA, Sigma-Aldrich, St. Louis, MO, USA, R8674). For the glycolytic stress assay, punches were incubated with 1 μM mix of rotenone/antimycin A mix and 50 mM 2-Deoxy-D-glucose (2-DG, Sigma-Aldrich, St. Louis, MO, USA, D6134). The assay program in the analyzer was then set as follows: 5 cycles of measurements for baseline, followed by the injection of the first drug, and then 4 cycles of measurements. Next, the second drug was injected, followed by another 4 cycles of measurements. Each cycle consisted of a mix (3 min), a wait (2 min), and a measure (3 min). After the assay, data were retrieved from the analyzer, data was collected, and statistical analysis was performed in Graphpad Prism 9 using one-way ANOVA, p-value < 0.05.

### Protein extraction and western blotting

Mechanically isolated retinas and RPE were lysed in RIPA buffer (ThermoScientific Chemicals, Waltham, MA, USA, J63324) containing a protease and phosphatase inhibitor cocktail I and II (Sigma-Aldrich, P8340, J63907, and J61022). Retinas were sonicated twice for 15 s while RPE was passed through a 27 1/2 G syringe needle, incubated on ice, and vortexed every 5 min for 20 min. Both lysates were centrifuged for 10 min at 14000 rpm at 4 °C, after which the supernatants were collected for western blotting. Protein quantification was performed using the Micro BCA Protein Assay Kit (ThermoFisher Scientific Inc, 23235), followed by protein (30 μg) separation in a 4–20% Novex Tricine SDS-PAGE (ThermoFisher Scientific Inc., XP04202) and transfer to PVDF membranes (Immobilon-FL; MilliporeSigma, Burlington, MA, USA, IPFL00010). Primary antibodies were incubated overnight at 4 °C, followed by washing and incubation with anti-mouse IRDye®680RD, anti-rabbit IRDye®680RD, and anti-mouse IRDye®800CW (all from LI-COR Biosciences, Lincoln, Nebraska, USA, 926-68070, 926-68071, 926- 32210). Membranes were incubated with the following primary antibodies: Total OXPHOS antibody cocktail (Abcam, Mouse, ab110413), HK I (Cell Signaling, Rabbit, 2024S), HK II (Cell Signaling, Rabbit, 2867S), PKM2 (Cell Signaling, Rabbit, 4053S), LDHA (Cell Signaling, Rabbit, 2012), EAAT3 (Thermo Fisher, Rabbit, 12686-1-AP). Immunoreactive signals were visualized using Oddessey CLx (LI-COR Biosciences) and the intensity signals were quantified using ImageJ software. Housekeeping proteins Actin (Cell Signaling, Mouse, 3700S) and GAPDH (Proteintech, Rabbit, 10494-1-AP) were used as internal control, and 3-month-old WT mice averages were used to calculate fold changes.

### Retinal immunohistochemistry

Enucleated eyes were fixed overnight at 4 °C by immersion in 4% paraformaldehyde made in D-PBS and sequentially infused with sucrose and Tissue-Tek O.C.T Compound (Sakura Finetek USA, Inc., Torrance, CA, 4583). Cryosections (8 μm) were cut on a cryostat HM 505E (Microm, Walldorf, Germany) equipped with a CryoJane Tape-Transfer system (Leica Inc., St. Louis, MO). For labeling, sections were washed with PBS, blocked in PBS supplemented with 1% BSA (PBS/BSA) and 0.1% Triton-X100 for 30 min, and incubated with primary followed by secondary antibodies conjugated with Alexa Fluor 488 and 594 (Molecular Probes, 1:1000) for 45 min at room temperature. The following primary antibodies were incubated with the retinas: GLUT-1 (Alpha Diagnostics, Rabbit, GT11-A), MCT1 & 3 (Rabbit, kindly donated by Dr. Nancy Philp). Nuclei were labeled with TO-PRO-3 (Thermo Fisher Scientific Inc, T3605). Sections were imaged using a laser scanning confocal microscope (Leica TCS-SP8, Leica, Exton, PA, USA) using the same acquisition parameters for each channel in the Leica confocal software LAS-X. Signal intensity was quantified within 200 μm of the optic nerve head, and the corrected total cell fluorescence (CTCF) was calculated for each area according to the following formula: CTCF = integrated density – (area of selected cell x mean fluorescence of background readings).

### Statistical Analysis

Data were analyzed using GraphPad Prism v10.1.1 (GraphPad Software) and are presented as the mean ± standard deviation (SD). Two-way ANOVA with Tukey’s Multiple Comparisons test (main row effect) and (simple effects within columns) and unpaired, two-tailed Student’s t-test were used to determine statistical significance between groups with an alpha value of 0.05. P values ≤ 0.05 were considered statistically significant.

## Author Contributions

J.S. and V.L.B. conceptualized and designed the study. K. J. and S. H. performed and helped analyze the Seahorse experiments. M. M. analyzed the immunohistochemistry data. J.S. wrote the manuscript, and all authors contributed to editing and finalization of the manuscript.

## Acknowledgements

The authors thank Dr. Anny Mulya and Dr. Alison Janocha for their technical assistance with the Seahorse experiments. The authors thank Dr. Nancy Philp (Thomas Jefferson University) for the MCT1 and MCT3 antibodies and for providing critical comments on this manuscript. The authors also thank David Schumick, BS, CMI for preparation of the illustration included in the manuscript.

## Data Availability Statement

Any information required to reanalyze the data reported in this paper is available from the lead contact upon request.

---

**FIGURE S3-1:**
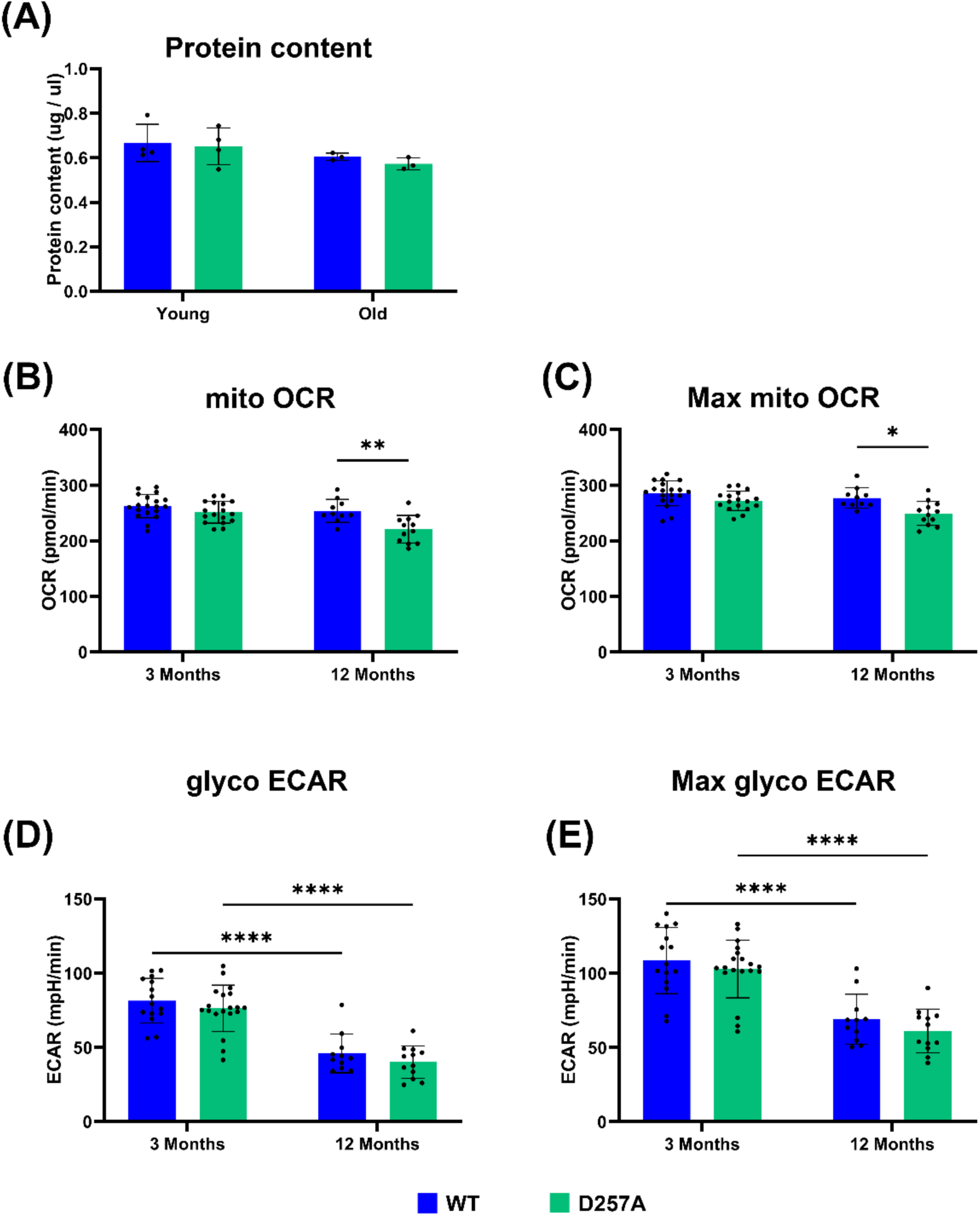
Seahorse analysis of ex vivo D257A retinal punches. **A)** Protein content measured by MicroBCA of young (3-month) and old (12-month) WT and D257A mice retina punches. **B)** Mitochondria associated basal oxygen consumption rate (OCR). **C)** Maximum mitochondria associated OCR after Bam15 injection. **D)** Basal glycolysis associated with extracellular acidification rate (ECAR). **E)** Maximum glycolysis associated with ECAR after rotenone/antimycin A injection. Data points = technical replicates / individual retina punches. Statistics: two-way ANOVA, p-value < 0.05.

**FIGURE S3-2:**
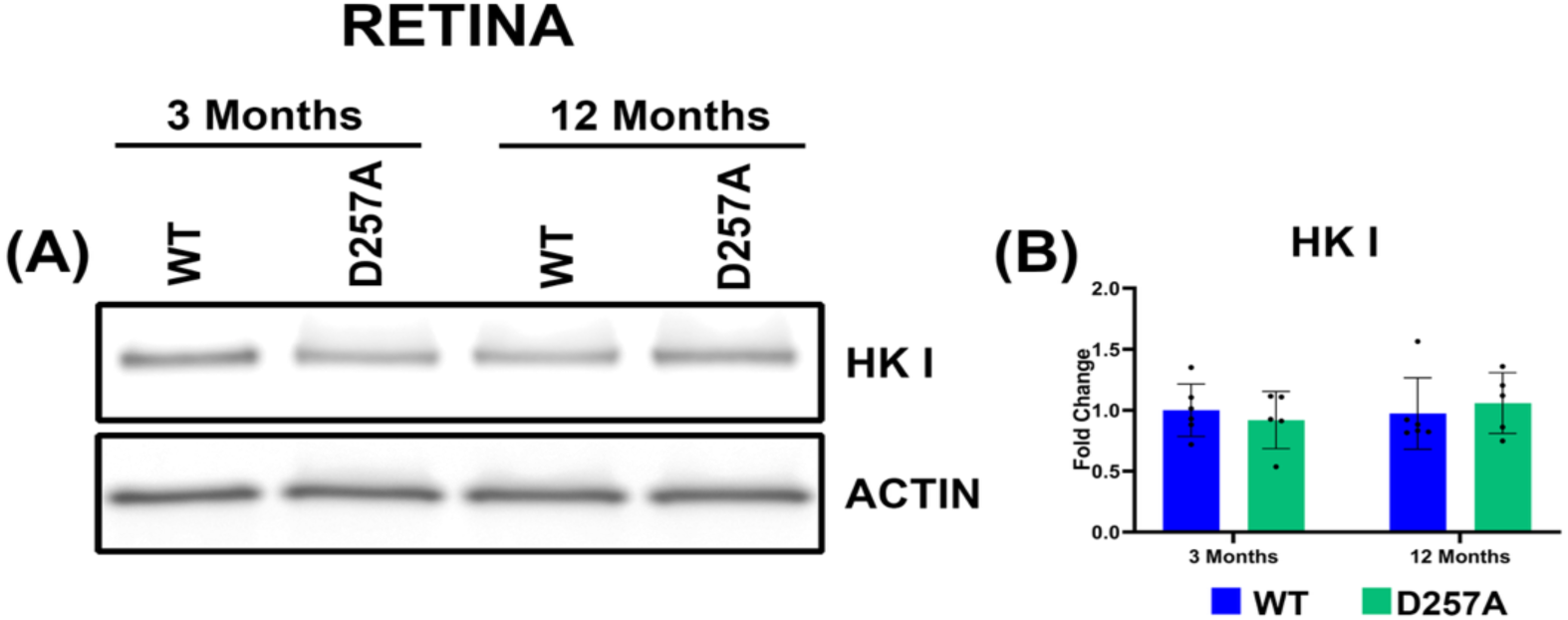
Alterations in key metabolic enzyme proteins in D257A retina. **A)** Representative immunoblot of Hexokinase I (HK I) in D257A retina. **B)** Quantification of HK I signal intensity in WT and D257A retina. Statistics: Data points = biological replicates (n=5); two-way ANOVA; p < 0.05.

**FIGURE S3-3:**
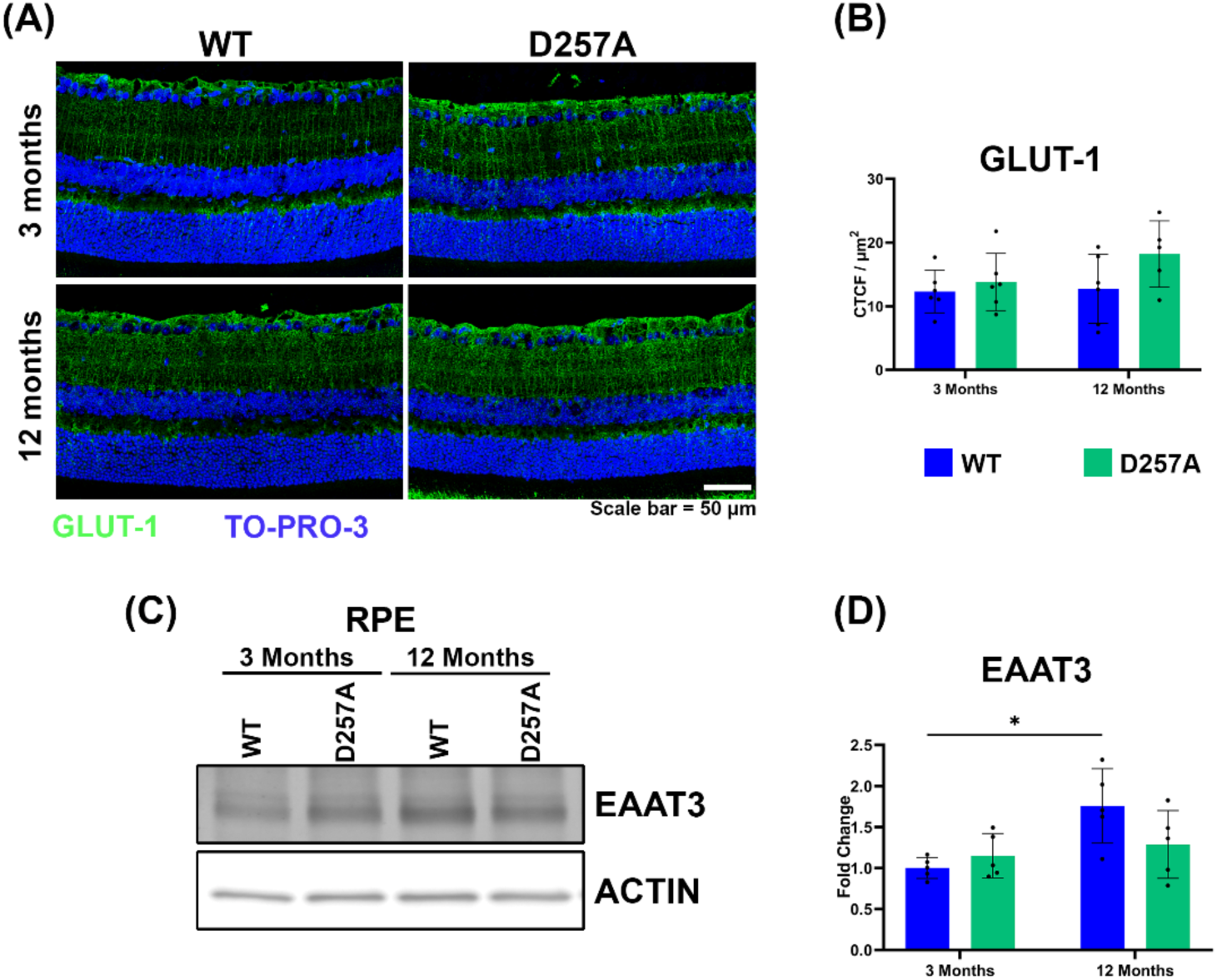
Investigation of transporters distribution and signal intensity in D257A retina and RPE. **A)** Representative confocal images of 3-month and 12-month-old retina GLUT1 staining. B) Quantification of retinal GLUT1 staining. C) Representative immunoblot of neuronal excitatory amino acid transporter (EAAT3) in 3-month and 12-month-old RPE. D) Quantification of EAAT3 signal intensity in WT and D257A RPE. Statistics: Data points = biological replicates (n=5); two-way ANOVA; p < 0.05.

